# Looking for Details: Fine-Grained Visual Search at the Foveal Scale

**DOI:** 10.1101/2025.11.09.687480

**Authors:** Sanjana Kapisthalam, Martina Poletti

## Abstract

Visual search has traditionally been conceptualized as the process by which the visual system identifies relevant objects in the extrafoveal field for subsequent foveation. Standard models implicitly assume there is no need for a finer-level search within the 1-degree foveola, as stimuli falling in this region are already viewed at high resolution. Using state-of-the-art high-precision eye tracking with arcminute level accuracy to localize the line of sight, we show that the visuomotor system is capable of implementing effective visual search strategies with striking precision within the foveated region. This active behavior, mediated by microsaccades, tiny saccades just a few arcminutes in size, allows for an efficient examination of the details falling in the central fovea. Our findings challenge the traditional visual search view and reveal that at each fixation a finer priority map of the foveal input is established, underscoring the dynamic nature of these representations.

## Introduction

Being able to actively scan the visual environment for a particular object (the target) among other stimuli (the distractors) is a fundamental skill for survival and something humans do daily; we regularly perform tasks of finding objects such as phones, wallets, keys or specific grocery items. Although these tasks are usually performed effortlessly, they are the result of complex processes weighing different aspects of visual stimuli, such as their perceptual salience and their degree of similarity to the target, to determine where to look next. To sample the visual environment, humans use saccadic eye movements. Saccades effectively implement visual search strategies, bringing objects of interest onto the center of gaze, where they can be processed at high resolution. It is well established that visual search is influenced by both top-down and bottom-up factors (Theeuwes, 1992, Theeuwes et al., 1998, Gaspelin et al., 2017, Gaspelin and Luck, 2019). Top-down factors refer to the influence of higher-level cognitive processes that guide attention and perception, such as attentional priorities, expectations, and task-specific goals (Baluch and Itti, 2011). Bottom-up factors, on the other hand, refer to low-level sensory cues that automatically capture attention, *i.e*., salient visual features such as color, motion, or abrupt changes in luminance (Connor et al., 2004). Both top-down and bottom-up factors interact dynamically during visual search, shaping how individuals explore and extract information from their visual environment (Dent, 2023, Wolfe, 2018, Theeuwes, 2018, Van Zoest and Donk, 2004).

According to the standard view, the visual system constructs a “priority map”, often described as a winner-take-all neural mechanism that directs both covert and overt attention (Bisley and Mirpour, 2019, Gregory J. Zelinsky and James W. Bisley, 2017, Mirpour et al., 2009, Desimone et al., 1995, Hobson et al., 1988). These maps are generally believed to represent the priority of objects in the extra-foveal visual field, where visual resolution is lower. To determine whether a high-priority object is indeed the desired target, foveation is typically required. Once the region containing the potential target is foveated, the 1-degree central foveal input is processed at high resolution. Consider the following scenario: You are looking for an item in a grocery store aisle located some distance away, where the region of interest occupies only a small portion of the visual field, which may be cluttered and rich in detail (Figure 1A). According to the traditional view, foveating this region allows the visual system to acquire a high-resolution snapshot of the central foveal input. Based on this detailed visual information, the observer can then determine whether the target is present. However, this perspective characterizes foveal vision during fixation as a largely passive process and overlooks the active role of the visuomotor system. Rather than simply taking a static snapshot of foveal input, the visuomotor system actively engages in small, precise eye movements. During fixation, both ocular drift and frequent microsaccades occur continuously (Poletti et al., 2020, Rucci and Poletti, 2015). Microsaccades, in particular, are precisely controlled and actively employed by the visuomotor system to explore the foveal input at a finer scale (Shelchkova et al., 2019, Intoy and Rucci, 2020). This raises the question of whether and how oculomotor behavior during fixation is engaged when we need to determine the presence of a target object in a complex and cluttered stimulus falling within the central foveal region.

**Figure 1.**
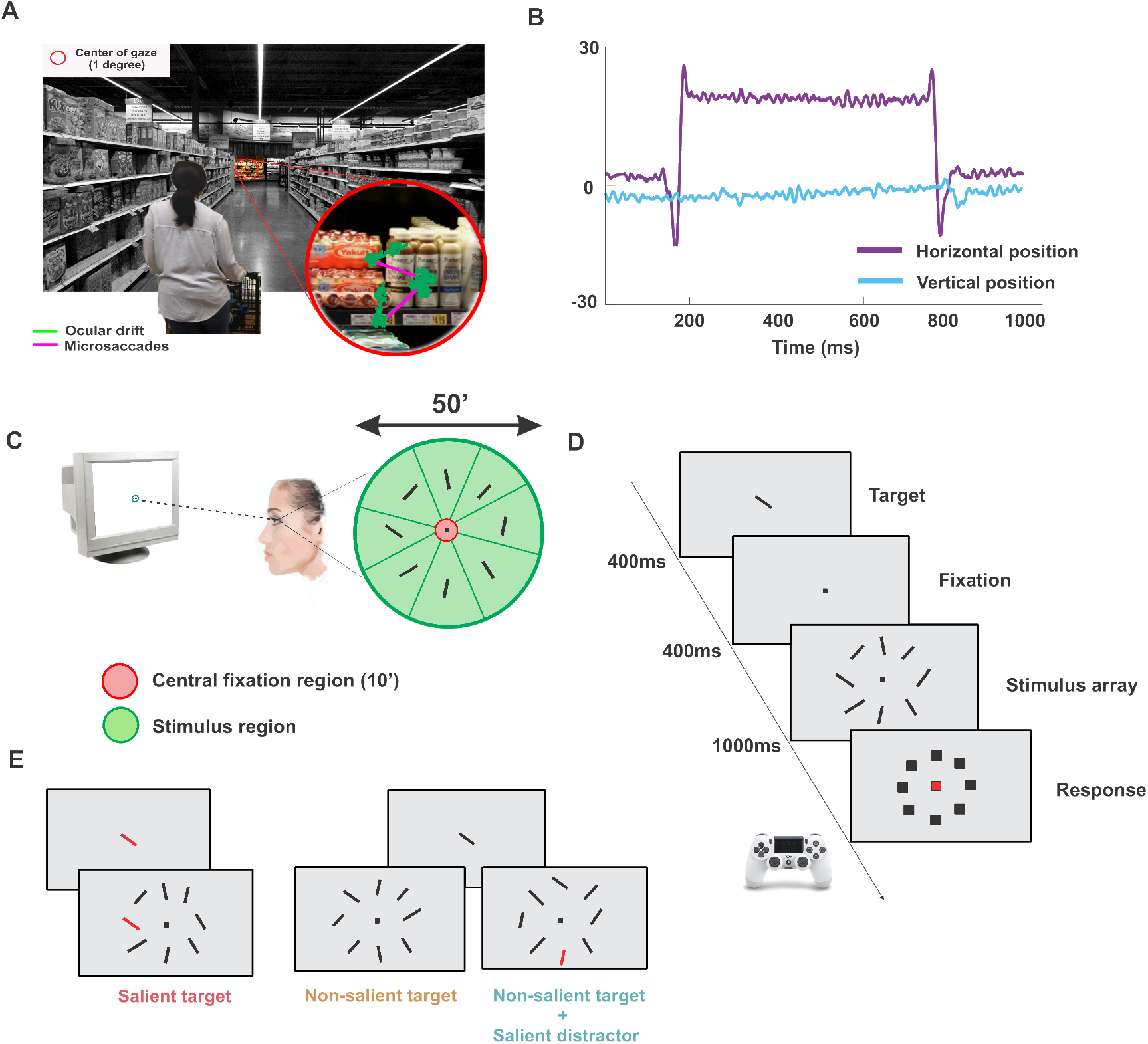
Experimental Paradigm. *A*, an example of an everyday situation in which foveal visual search could occur. The inset shows an enlargement of the visual input to the central fovea with an example of microsaccades and ocular drift overlayed on the stimulus. *B*, an example eye movement trace from the task. *C*, spatial arrangement of the stimuli in the task. Stimuli were presented foveally (*i.e*., spanning a region of less than 1^*◦*^ in diameter at the center of gaze). The target and the stimuli in the array consisted of small bars (6^*′*^*×*2^*′*^) each tilted of a different angle. The illustration also shows how gaze position was classified based on its location on the stimulus array. Gaze position outside the green contour was classified as on background. *D*, time course of a typical trial. After a brief fixation period, in which the target stimulus was presented at the center of the display, an array with 8 stimuli was presented for 1s. Subjects were then asked to determine whether or not the target was in the array and in which of the eight locations it was presented. *E*, experimental conditions. Experimental conditions were defined based on whether or not the target was salient and whether or not a salient distractor was present. Perceptually salient items were distinguished by a unique color that set them apart from other objects in the array.

Addressing this question is challenging because, in the foveola, target and distractors are only a few arcminutes away from each other. As a result, high-precision eye-tracking and accurate localization of the line of sight at the arc-minute level are needed. This cannot be achieved with commercial video-based eyetrackers, which yield an error in gaze localization that can be as large as 1 degree (Holmqvist and Blignaut, 2020, Holmqvist and Andersson, 2017). To overcome this limitation, here we relied on a unique blend of tools by coupling a high-precision, custom-made Digital Dual Purkinje Image Eye-tracker (Wu et al., 2023) with a system for gaze-contingent display control, which allows for a more accurate localization of the line of sight compared to standard methods (Poletti and Rucci, 2016, Santini et al., 2007). While monitoring fixational eye movements, we asked subjects to determine whether a target detail was present in the foveal input. Our work shows that the visuomotor system is capable of establishing fine-grain priority maps of the 1-degree visual input at the center of gaze to effectively drive visual exploration of complex foveal stimuli.

## Results

We examined how oculomotor behavior at fixation is modulated, and the factors driving microsaccades when observers are required to localize a target in a complex foveal stimulus. Participants were asked to determine the location of a target among an array of similar items. Stimuli consisted of small bars (6^*′*^*×*2^*′*^). The stimulus array covered an area of 0.5^*◦*^ in diameter, approximately half the size of the foveola (figure 1C). Participants were first presented with a target, a tilted bar. After a blank period, the stimulus array was displayed for 1 second. At the end of the trial, subjects were required to indicate the location of the target in the array (figure 1D). The saliency of the stimuli in the array was adjusted by changing their color, whereas task relevance was adjusted by modulating the degree of tilt of each bar. Stimuli with an orientation similar (*±*10^*◦*^) to the target were considered task-relevant. We distinguished between three main conditions based on salience and relevance of the stimuli and the target: (*a*) salient target with non-salient distractors, (*b*) non-salient target with non-salient distractors, and (*c*) non-salient target with a salient distractor in the array (figure 1E). Each of these conditions had the same probability of occurrence, and they were interleaved during the task. The degree of tilt of the target and the other stimuli in the array was determined individually to achieve an overall performance of approximately 75% correct responses across all three conditions. Subjects were not forced to maintain fixation at the center of the array when stimuli were presented, and they could freely move their eyes during the trial.

Oculomotor behavior was monitored using a high-precision eye tracker (Wu et al., 2023) (see Methods for details). Figure 2A illustrates the average 2D normalized probability of gaze position over time with respect to the location of the target. When the stimulus array was initially presented, observers’ gaze position was around the center of the display, so every item in the array was approximately 15^*′*^ away (center-to-center distance) from the preferred locus of fixation. However, subjects did not maintain their gaze around the center throughout the trial. Instead, over the course of the trial, gaze position generally moved toward one of the items in the array, usually the target (figure 2A,B). When the target was salient, participants’ accuracy in identifying the target was high (92%±2%) (figure 4B) and gaze position was tightly clustered around the target as early as 300 ms from stimulus onset (first column in figure 2B). We then determined how gaze distribution evolved over the course of a trial relative to the salient target location (in figure 2B shown at location 0, see also Movie S1). Early on (0-300 ms from stimulus onset) gaze remained near the central marker, and gaze position was uniformly distributed across all the stimuli in the array (F(7,56) = 0.88, p = 0.52,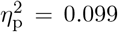), whereas, between 300-500 ms, and in the subsequent time intervals, the probability of gaze being on the target was higher than the probability of being on every other stimulus, indicating an early orienting toward the salient target (*F* (7, 56) = 77.35, Greenhouse-Geisser corrected *p <* 0.001, 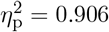. Tukey’s HSD post hoc comparisons p ¡ 0.001 for all locations vs. the target location for the 300-500 ms interval. 500-700 ms interval: *F* (7, 56) = 128.23 Greenhouse-Geisser corrected *p <* 0.001, 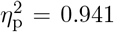. 700-1000 ms interval: *F* (7, 56) = 161.62, Greenhouse-Geisser corrected *p <* 0.001, 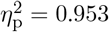).

**Figure 2.**
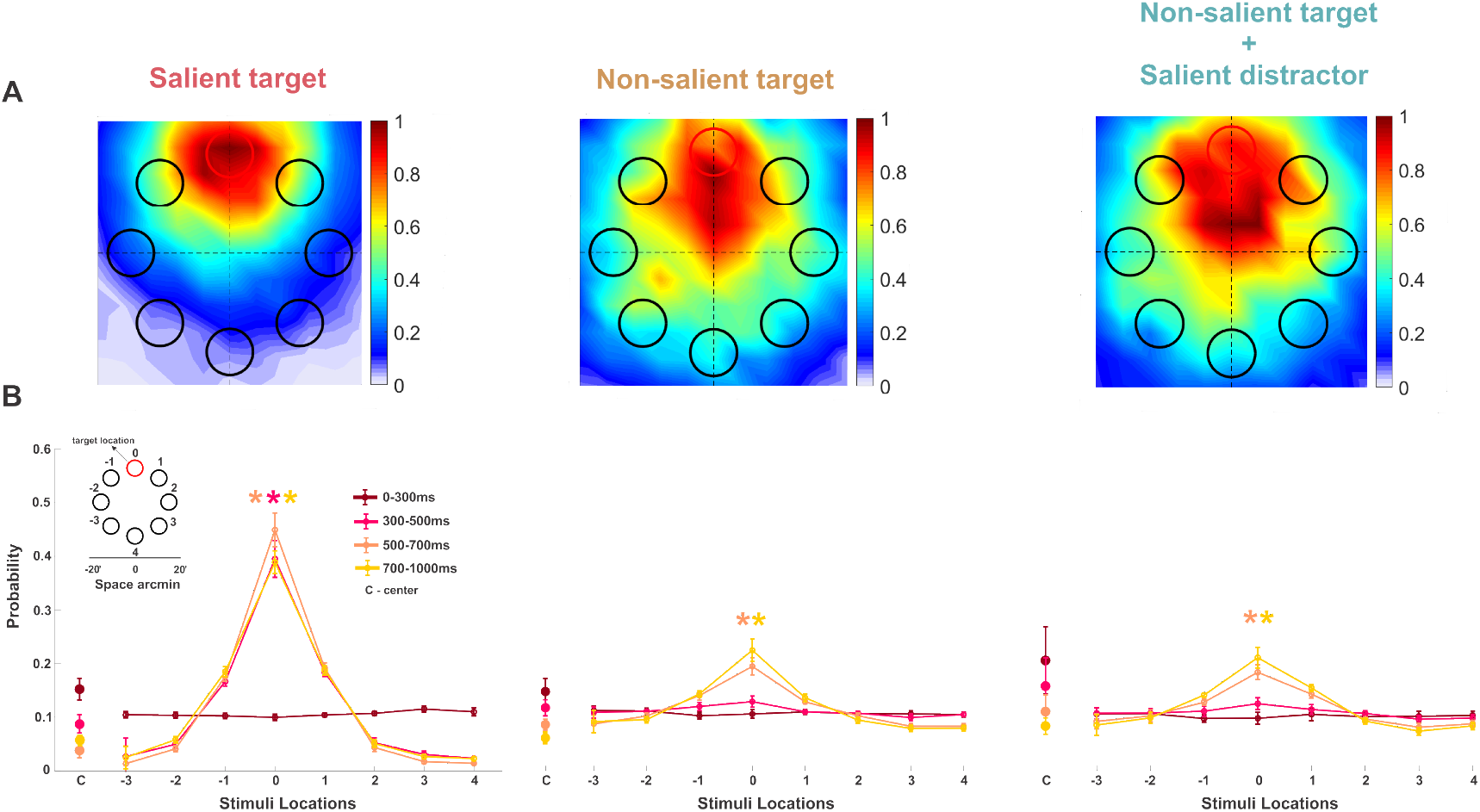
Gaze distribution in the task. *A*, probability of gaze position being over different stimuli and in the center (C) of the array at different time intervals during the trial. The shaded gray regions highlight the statistically significant comparisons. The inset shows the stimuli’s spatial arrangement with respect to the target location. Asterisks mark a statistically significant difference (p*<*0.001, repeated measures ANOVA). Error bars are SEM. *B*, average 2D normalized probability distributions of gaze position during the last 700 ms–1000 ms of the stimulus presentation over the stimulus array with respect to the target location (red circle on top).

In contrast, when the target was not salient, gaze position was broadly distributed across the stimulus array, and clustered toward the target only at a later time (Figure 2B, middle column). Between 0-500 ms, gaze was uniformly distributed across all stimuli locations (from 0-300 ms: *F* (7, 56) = 0.48, *p* = 0.75, 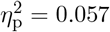; from 300-500 ms: *F* (7, 56) = 2.40, *p* = 0.108, 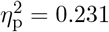). Only after 500 ms, the probability of gaze being on the target was higher than the probability of being on every other stimulus in the array (from 500-700 ms: *F* (7, 56) = 20.76, Greenhouse-Geisser corrected *p* = 7.53 *×* 10^*−*5^, 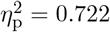, and from 700-1000 ms: *F* (7, 56) = 25.16, Greenhouse-Geisser corrected *p* = 3.09 *×* 10^*−*5^, 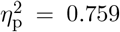, all post hoc comparisons between the target and all other locations were statistically significant, *p <* 0.001). Interestingly, this behavior did not change when a salient distractor was present in the stimulus array (Figure 2B, right column; see also Movie S1) (from 500-700 ms: *F* (7, 56) = 18.21, Greenhouse-Geisser corrected *p* = 4.34 *×* 10^*−*14^, 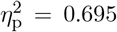, and 700-1000 ms: *F* (7, 56) = 22.09, Greenhouse-Geisser corrected *p* = 1.93 *×* 10^*−*4^, 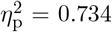, for all post hoc comparisons between the target and all other locations *p <* 0.001).

During the last 700 to 1000 ms of stimulus presentation, the average gaze position was 4.5^*′*^±2^*′*^ away from the salient target. However, this distance doubled when the target was not salient (figure 3A). A repeated measures ANOVA revealed a significant main effect of condition on gaze distance (F(2,16) = 33.11, Greenhouse-Geisser corrected p = 0.00005, 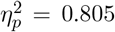). Post hoc comparisons confirmed that gaze was closer to the target when it was salient, than when it was not salient (p = 0.002 and p = 0.001, without and with a salient distractor respectively). These results illustrate that if a stimulus is salient and relevant, it influences fixation behavior at the foveal scale. However, if the stimulus is a salient singleton, but is irrelevant to the task and acts as a distractor, neither gaze distribution nor performance is affected.

**Figure 3.**
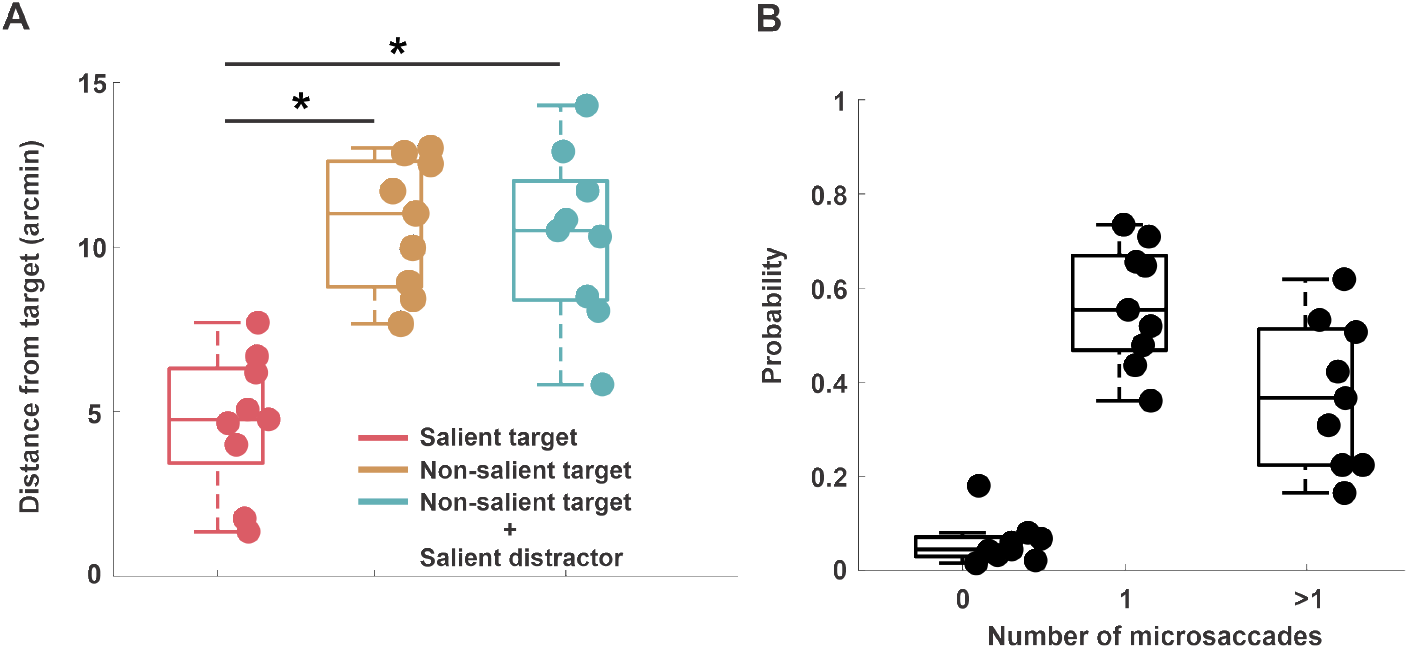
Average gaze position distance and average rate of microsaccades. *A*, Euclidean distance of gaze position from the target between 700 ms - 1000 ms. Asterisks mark a statistically significant difference (*p <* 0.001, repeated measures ANOVA). *B*, average rate of microsaccades in the task. In both panels, the center line of each box represents the median, and the upper and lower box boundaries indicate the 75th and 25th percentiles, respectively. Individual markers represent individual subject data.

### Influence of saliency on microsaccadic behavior

The changes in gaze position described above were driven by microsaccades. Most trials had at least one microsaccade (93%±4%) and in 37%±15% of the trials, there were two or more microsaccades (figure 3B). The average amplitude of microsaccades (16.9^*′*^±2^*′*^) approximately matched the target eccentricity (inset figure 4C). We then examined how the latency of microsaccades was influenced by saliency. Our results show that it took 311 ms *±* 30 ms for the visuomotor system to execute a microsaccade toward a salient target. But this time increased by 121 ms *±* 81 ms when the target was not perceptually salient (two-tailed paired t-test, t(8) = 4.4, p = 0.002, Cohen’s d = 1.78) (figure 4A). This behavior was also accompanied by a higher performance in the salient target condition compared to the other conditions (salient vs. non-salient target: 92%±2% vs. 63%±7%;two-tailed paired t-test, t(8) = −9.7, p *<* 0.00001, Cohen’s d = 4.3) (figure 4B). On the other hand, both latency (two-tailed paired t-test, t(8) = −1.04, p = 0.32, Cohen’s d = 0.06) and performance (two-tailed paired t-test, t(8) = −1.7, p = 0.12, Cohen’s d = −0.33) were unaffected by the presence of a salient distractor (figure 4C,D). This finding illustrates that saliency influences the latency of microsaccades only when it belongs to the target, *i.e*., the latency of microsaccades was lower when microsaccades were directed towards a salient target. Notably, we observed similar latency differences between conditions even for large saccades directed toward stimuli presented extrafoveally (see Supplementary Material, figure S1). This suggests that comparable visual search mechanisms operate both within the fovea and extrafoveally.

**Figure 4.**
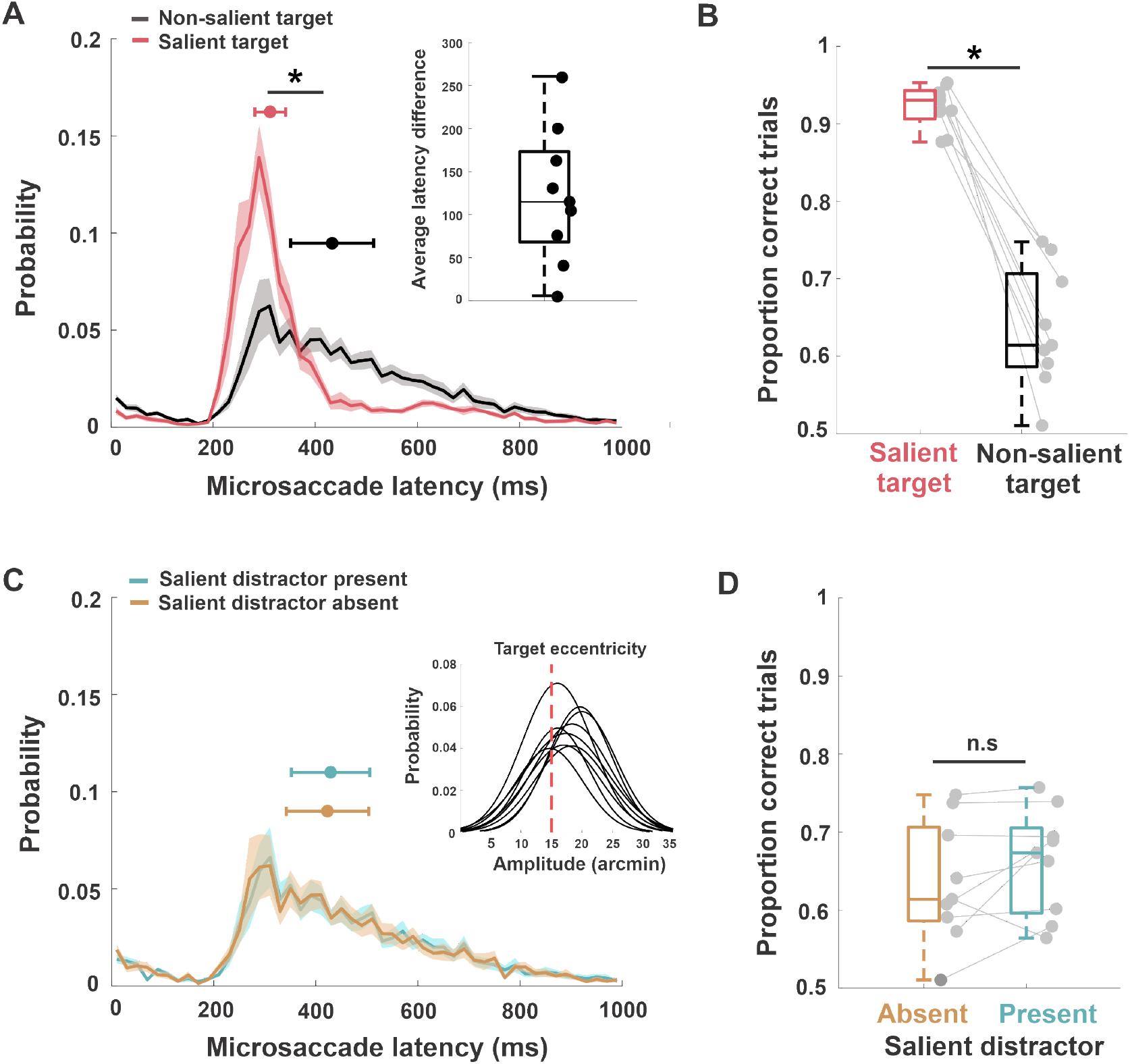
Microsaccade latency and response accuracy as a function of target saliency. *A*, average latency distributions of the first microsaccade for a salient and non-salient target. Inset—average latency difference between the two conditions. *B*, average proportion of trials in which participants correctly identified the target location, both in the presence of a salient and non-salient target. The asterisk marks a statistically significant difference (*p <* 0.01, two-tailed paired t-test). *C*, average latency distributions of the first microsaccade in the presence and in the absence of a salient distractor. Inset - microsaccade amplitude for individual subjects in the task across all trial types. The red dashed line shows the eccentricity of the target from the center of the display. *D*, average proportion of trials in which participants correctly identified the non-salient target location, both in the presence and absence of a salient distractor. Shaded error bars in *A* and *C* are SEM, and error bars on top mark the median latency and SEM across subjects. In *B* and *D*, the center line represents the median, and the upper and lower box boundaries indicate the 75th and 25th percentiles, respectively. Filled circles without error bars represent individual subjects.

To further assess the impact of saliency on oculomotor behavior, we examined the accuracy of microsaccades (*i.e*., the probability of the first microsaccade landing on the target) (see Methods and figure 1B for details). When the target was salient, microsaccades were more accurate (figure 5A): 50%±15% of the first performed microsaccades landed on the salient target, compared to 28%±5% when the target was non-salient and no salient distractor was present, and 29%±6% when a salient distractor was present. A repeated-measures ANOVA revealed a significant main effect of condition on microsaccade accuracy (F(2,16) = 20.81, Greenhouse-Geisser corrected p = 0.0014, 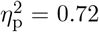). Post hoc pairwise comparisons showed that microsaccade accuracy was higher when the target was salient compared to when it was not (salient distractor absent: p = 0.0043; present: p = 0.0044). There was no significant difference between the two non-salient target conditions (p = 0.51). These results demonstrate that saliency enhances microsaccade accuracy, and that the presence of a salient distractor does not reduce the proportion of target-directed microsaccades.

**Figure 5.**
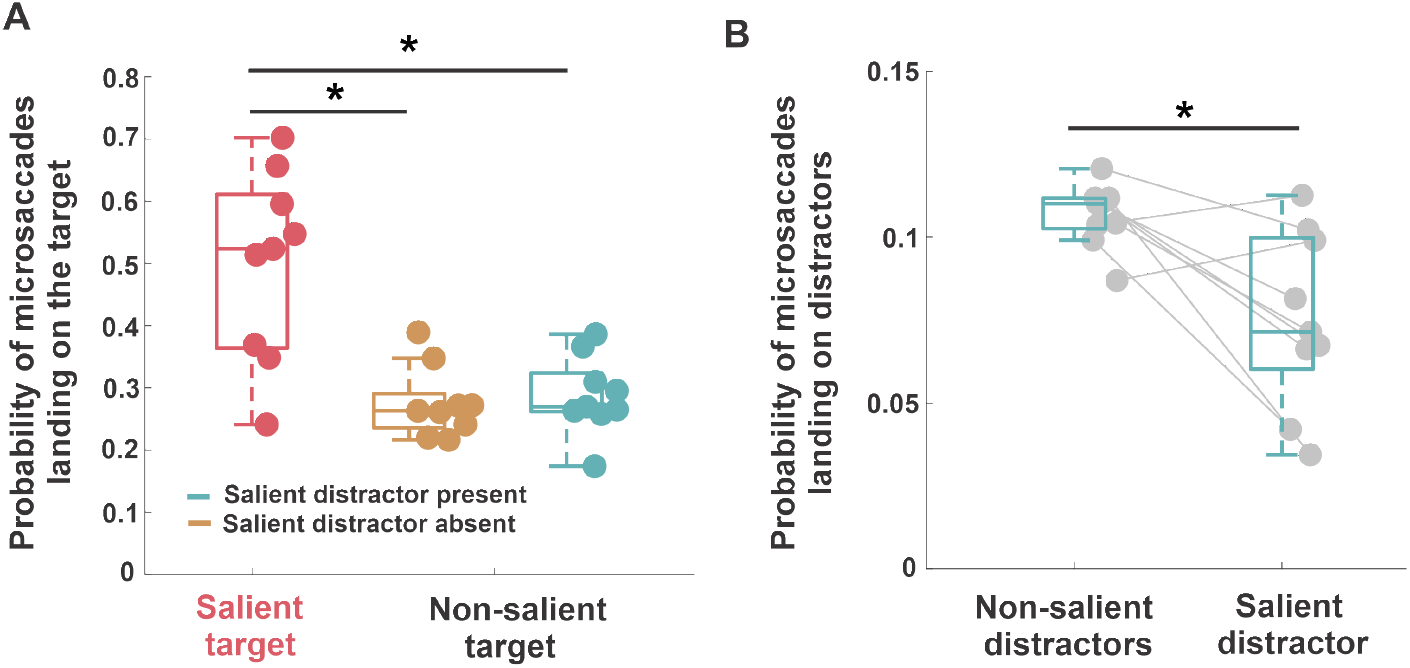
Microsaccade accuracy as a function of saliency. *A*, probability of the first microsaccade landing on target in different conditions. Asterisk marks a statistically significant difference (p=0.001, repeated-measures ANOVA). *B*, probability of the first microsaccade landing on one of the distractors when the target was non-salient. Asterisk marks a statistically significant difference (p=0.01, two-tailed paired t-test). In both panels, the center line of each box represents the median, and the upper and lower box boundaries indicate the 75th and 25th percentiles, respectively. Filled circles without error bars represent individual subjects.

Further, our results suggest that microsaccades towards salient distractors were suppressed (figure 5B); the probability of microsaccade landing on a salient distractor was slightly lower compared to the probability of landing on a non-salient distractor (8%±2% vs. 11%±0.9%; two-tailed paired t-test, t(8) = 3.2, p = 0.01, Cohen’s d = 1.4), possibly indicating avoidance of salient distractors. Interestingly, when the target was not salient and a salient distractor was absent, the probability of microsaccades landing on the target in the correct trials was higher than in incorrect trials (figure S2A, 32%±9% vs. 19%±3%; two-taile paired t-test, t(8) = 3.2, p = 0.01, Cohen’s d = 1.64). We observed a similar behavior even when a salient distractor was present (figure S2B, 33%±8% vs. 20%±5%; two-tailed paired t-test, t(8) = 4.7, p = 0.001, Cohen’s d = 1.73), indicating that accurate microsaccades were associated with better localization performance.

### Influence of task-relevance on microsaccadic behavior

To examine the influence of task-relevance on microsaccades, trials were classified as either having high or low task-relevance; when the orientation of one item in the array was within *±*10^*◦*^ of the target’s orientation, trials were labeled as trials with a high task-relevance distractor (inset figure 6) otherwise they were categorized as trials with low task-relevance distractors (see inset in figure 6 purple shaded region). Figure 6 illustrates that in the presence of a high task-relevant distractor, the probability of a microsaccade landing on the target dropped (high task-relevant distractor absent vs. present: 41%±8% vs. 20%±11%; two-tailed paired t-test, t(8) = 4.02, p = 0.003, Cohen’s d = 1.88), indicating that the similarity of a stimulus to the target is a factor that directs microsaccades in a visual search task. Therefore, microsaccades were influenced both by the saliency and by the task relevance of distractors.

**Figure 6.**
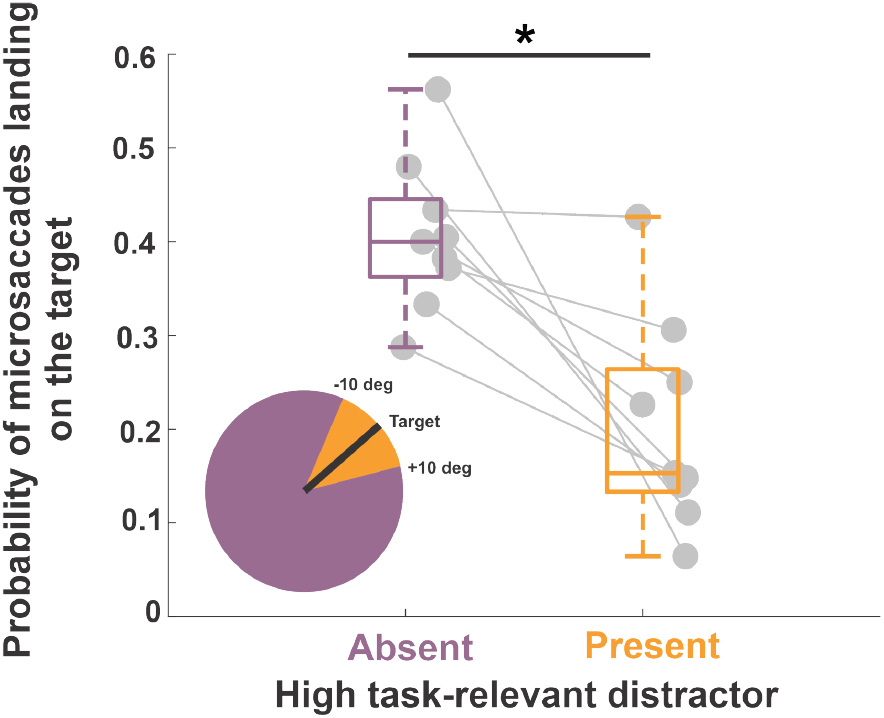
Microsaccade accuracy as a function of task-relevance. Average probability of the first microsaccades landing on the target in trials with low and high task-relevance distractors respectively. In low task-relevance distractors trials the orientation of all non-target items is more than 10^*◦*^ away from that of the target. In high task-relevance distractor trials, the orientation of one item is less than 10^*◦*^ away from that of the target orientation. Asterisks mark a statistically significant difference (two-tailed paired t-test, p=0.003). The center line of the box plot represents the median, and the upper and lower box boundaries indicate the 75th and 25th percentiles, respectively. Filled circles without error bars represent individual subjects.

## Discussion

Because high-resolution vision is limited to the central 1^*◦*^ foveola, and humans rely on saccades to bring objects of interest into foveal view, visual search has naturally been studied in the context of exploring scenes that span many degrees of visual angle(Theeuwes, 1992, Theeuwes et al., 1998, Casteau and Smith, 2020, Theeuwes, 2019, Gaspelin et al., 2015, Gaspelin and Luck, 2018, Gaspelin et al., 2019). It is generally assumed that active visual search is only needed in the presence of extra-foveal stimuli. However, even during short foveation periods, our eyes continue to move with tiny saccades less than half a degree in size, known as microsaccades (Rucci et al., 2007, Rucci and Poletti, 2015, Poletti, 2023). Whereas, microsaccades for a long time were mostly considered random movements to prevent image fading (Martinez-Conde et al., 2006, McCamy et al., 2012), it is now clear that they serve to precisely reposition the preferred locus of fixation during tasks requiring fine spatial vision (Ko et al., 2010, Poletti et al., 2017, Shelchkova et al., 2019). This behavior is not a mere reflex to maintain the fixated object at the preferred locus of fixation; on the contrary, it is modulated by task requirements and by cognitive factors (Shelchkova et al., 2019, Intoy and Rucci, 2020). This raises the possibility that visual search may not be limited to the extra-foveal visual field. Rather, finer search strategies might be implemented when examining the details falling in the foveola. Here, we address this issue and examine the spatio-temporal dynamics of microsaccades using custom-made high-precision eye-tracking coupled with a gaze-contingent display system to achieve higher accuracy in gaze localization (Santini et al., 2007, Wu et al., 2023).

In our study, observers were asked to identify a target in a cluttered foveal stimulus, covering less than 1 deg of visual angle. Importantly, as in normal viewing conditions, subjects freely moved their eyes during stimulus presentation, and their oculomotor behavior in the task was spontaneous, *i.e*., fixation on a point was not enforced. Our findings show that subjects engage in active visual search at the foveal scale using spontaneous microsaccades as small as 10^*′*^ (0.16^*◦*^). Microsaccade precision and reaction times were modulated both by salience and task relevance of the different details falling in the foveola. Subjects consistently prioritized task-relevant locations within the foveal input and appeared to actively suppress salient distractors. These findings provide crucial evidence further supporting the idea that fine oculomotor behavior during fixation is not solely driven by bottom-up re-centering mechanisms but is also the result of active, top-down-driven visual scanning strategies (Shelchkova et al., 2019). Further, they indicate that the visual system has access to a fine-grain saliency and priority map representation of the stimuli at the foveal scale.

These results inevitably raise the question of why humans need a fine-grain priority map of the central foveal input. It has been shown that fine spatial vision is not uniform in the foveola (Poletti et al., 2013, Intoy et al., 2021). Moreover, visual acuity at this scale is also known to be influenced by factors such as crowding (Coates et al., 2018, Clark et al., 2024, Shamsi et al., 2022, Bondarko et al., 2024), wherein nearby stimuli interfere with the perception of the target, which reduces acuity in normal viewing conditions when the input to the foveola is complex and rich in detail. Ultimately, microsaccades compensate for this non-uniform foveal vision by bringing details of interest closer to the preferred locus of fixation to be viewed at an even higher resolution (Shelchkova et al., 2019). Therefore, a fine-grain priority map of the foveal input may be essential for effectively identifying relevant stimuli at this scale and to enable finer visual examination.

As noted previously (Veale et al., 2017), there is a need for biologically plausible saliency and priority map models. Traditional models assume the existence of a static priority/saliency reference map of the extra-foveal space that is used to guide visual exploration across the scene (Itti and Koch, 2000, 2001, Foulsham and Underwood, 2008, Hayes and Henderson, 2021). However, saliency and priority are not fixed qualities of the internal representation of the scene; rather, they are continuously modulated not only by changes in the external stimulus but also by the ongoing oculomotor behavior (Hafed, 2013, Hafed et al., 2015). Besides the need to account for the dynamic nature of priority/saliency representations, our findings indicate that current models should also account for the existence of fine-grain priority maps of the foveal input. At each fixation, the visual system not only assesses and possibly updates extra-foveal priority/saliency maps, which are used as a reference to guide saccades, but it also generates afresh a finer scale map of the foveal input. If this map highlights details of interest, the visuomotor system engages in a finer exploration. Otherwise, a saccade is planned towards items of interest in the visual periphery. Ultimately, the content of this map and its interplay with the extra-foveal priority/saliency reference map determine whether or not fixation is maintained. Hence, any model of priority/saliency needs to incorporate not only this fine-grain map of the foveal input but also its dynamic nature.

While our study sheds light on the active visual search strategies employed at the foveal scale, several intriguing questions remain unanswered. Priority maps of the extra-foveal space have been shown to guide not only saccades but also reaching (Bisley and Mirpour, 2019, Bisley and Goldberg, 2010). This raises the question of whether the fine-grain priority map of the central 1^*◦*^ region of the visual field is exclusively used to guide microsaccades or if it can be accessed by systems in other sensory modalities. This fine-grain map could be useful for tasks requiring fine visuomotor coordination, such as threading a needle, grooming animals, precision shooting, or during fine surgical procedures, etc. Further research is necessary to address these questions.

In summary, leveraging high-resolution eye tracking and accurate localization of the line of sight, we show that humans use microsaccades as small as 10^*′*^ in size to explore task-relevant details in the foveal input. Overall, our results indicate that humans are capable of establishing high-resolution priority maps of the 1^*◦*^ visual input at the center of gaze, which are used to effectively guide visual exploration at the foveal scale.

## Methods and Procedures

### Observers

Nine participants (4 males, age range 18-27) took part in the main experiment. Participants were students from the University of Rochester with uncorrected 20/20 vision. Eight subjects were naïve and one subject was one of the authors. Four subjects (2 males) participated in the extra-foveal search task, a control experiment in which stimuli were presented extrafoveally. Two of these subjects also participated in the main experiment. This research was approved by the University of Rochester’s Institutional Review Board (IRB) for human subjects research. Subjects were invited for an initial screening session involving a Snellen eye-chart test. For subjects able to achieve 20/20 vision in the right eye without glasses or contact lenses, this initial screening was followed by a detailed review of the consent form. Informed consent was obtained and documented only after the subject understood the information on the consent form, consented, and signed to participate in the study.

### Experimental Setup

Stimuli were displayed on an ASUS monitor at a refresh rate of 144 Hz and spatial resolution of 1,920 × 1,080 pixels (1 pixel = 0.53^*′*^). Observers performed the task monocularly with their right eye while the left eye was patched. A dental-imprint bite bar and a headrest were used to prevent head movements. The movements of the right eye were measured by means of a custom-built digital Dual Purkinje Image (dDPI) eye tracker, a system with an internal noise of approximately 20” and a resolution of 1 arc-minute (Wu et al., 2023). Vertical and horizontal eye positions were sampled at 1 kHz and recorded for subsequent analysis. Stimuli were rendered by means of EyeRIS (Santini et al., 2007), a custom-developed system allowing for flexible gaze-contingent display control. This system acquires eye movement signals from the eye tracker, processes them in real time, and updates the stimulus on the display according to the desired combination of estimated oculomotor variables.

### Eyetracking Calibration Procedure

Every session started with preliminary setup operations that lasted a few minutes. The subject was positioned optimally and comfortably in the apparatus. Subsequently, a calibration procedure was performed in two phases. In the first phase, subjects sequentially fixated on each of the nine points of a 3 × 3 grid, as is customary in oculomotor experiments. These points were located 1.32^*◦*^ apart on the horizontal and vertical axes. In the second phase, subjects confirmed or refined the mapping given by the automatic calibration. In this phase, they fixated again on each of the nine points of the grid while the location of the line of sight, estimated on the basis of the automatic calibration, was displayed in real time on the screen. Subjects used a controller to correct the predicted gaze location, if necessary. The calibration mapping was then updated based on these corrections. This dual-step calibration allows for a more accurate localization of gaze position than standard single-step procedures, improving 2D localization of the line of sight by approximately one order of magnitude (Poletti and Rucci, 2016, Poletti et al., 2013). The manual calibration procedure was repeated for the central position before each trial to compensate for possible microscopic head movements that may occur even on a bite bar.

### Experimental paradigm

Subjects were briefly shown a target (a 6^*′*^*×*2^*′*^ tilted bar), then, after a short delay, a circular array of stimuli similar to the target but with different orientations was presented. The stimulus array was contained within the central half-degree region of the display where subjects were already fixating (figure 1C) and was presented for 1s. At the end of the trial, subjects were asked to determine whether the target was present in the array and, if so, to identify its position in the array by moving a cursor to the location of the target (figure 1D). If subjects could not locate the target they could report it as missing. The degree of tilt of the target in the array was tailored individually to attain an overall performance of approximately 75% correct responses across all three conditions. Subjects were allowed to freely look at the stimulus and their oculomotor behavior in the task was spontaneous. The target was present in 95% of trials across all conditions with an overall average miss rate of 2%±1%. In the remaining 5% of trials, the target was absent. In these trials, on average, the correct rejection rate was of 66%±6% and false alarm rate was 33%±6%.

Stimuli salience and task relevance were systematically varied in the experiment. The target could be either perceptually salient (*i.e*., different color from the rest of the stimuli in the array) or non-salient (figure 1E). In the latter case, salient distractors could be either absent or present. Task relevance was modulated by changing the degree of tilt of the distractors. Stimuli with an orientation similar (*±*10^*◦*^) to the target’s orientation were considered task-relevant (*e.g*., with a 135^*◦*^ tilted target a 125^*◦*^ tilted distractor has high task-relevance, and a 45^*◦*^ tilted distractor has low task-relevance). All conditions were interleaved.

### Analysis of behavioral data and statistics

Recorded eye movement traces were segmented into separate periods of drift and saccades. Classification of eye movements was performed automatically and then validated thoroughly by the authors. Periods of blinks were automatically detected by the digital DPI eye tracker and removed from data analysis. Only trials with optimal, uninterrupted tracking were selected for data analysis. Eye movements with minimal amplitude of 3^*′*^ and peak velocity higher than 3^*◦*^/s were selected as saccadic events. Saccades with amplitudes less than 0.5^*◦*^ were defined as microsaccades. Periods that were not classified as saccades or blinks were labeled as drifts.

To assess microsaccade accuracy, we classified microsaccades based on their landing positions. Microsaccades that landed within a circular region with a 10^*′*^ diameter centered on the fixation marker were classified as targeting the center of the stimulus array (red-shaded area in figure 1C). On average, these comprised 3%*±*0.8% of all microsaccades. Those landing outside a 50^*′*^ diameter region around the center were classified as targeting the background, accounting for 5%*±*2% of microsaccades on average. For all remaining microsaccades, we computed the Euclidean distance between each landing position and the items in the array to determine the closest stimulus target (green-shaded area in figure 1C).

All statistical analyses were conducted using the MATLAB Statistical Toolbox. We assessed sphericity assumption in the repeated-measures ANOVAs using the Maulchy test. When the sphericity assumption was violated (p *<* 0.05), Greenhouse-Geisser corrected p-values are reported; otherwise, uncorrected p-values are presented. All post hoc comparisons were adjusted using the Tukey-Kramer correction.

## Acknowledgements

This work was funded by Meta Reality Labs and NIH EY001319 grant to the Center for Visual Science at the University of Rochester. We thank Claire Corbeaux for their input while writing the paper.

