## Supplementary material for "Looking for Details: Fine-Grained Visual Search at the Foveal Scale"

**Movie S1**: Average normalized 2D probability distribution of gaze position relative to the target location (red circle) across the time course of a trial, shown separately for each condition.

**Foveal vs. extrafoveal visual search:** Our results show that visual search is also carried out at the foveal scale as an active process that relies on microsaccades to locate a target amongst an array of similar-looking details. This finding raises the question of how comparable visual search strategies at the foveal scale are with search strategies enacted when subjects view larger visual scenes. To address this question, we replicated our foveal search task extrafoveally at $5^o$ eccentricity with 4 subjects. This is a typical search task and closely resembles those used in previous research (Theeuwes, 1992, Theeuwes et al., 1998, Casteau and Smith, 2020, Theeuwes, 2019, Gaspelin et al., 2015, Gaspelin and Luck, 2018, Gaspelin et al., 2019). Stimuli size in extrafoveal conditions was adjusted accordingly to account for cortical magnification(Rovamo and Virsu, 1979).

The effect of visual saliency of the target on saccade latencies was also present for larger saccades; latencies were shorter when the target was visually salient (salient vs. non-salient target: 232ms$\pm$30ms vs 444ms$\pm$93ms),and each individual subject showed a significant difference in microsaccade latency between salient and non-salient target conditions (Wilcoxon rank-sum test: all $p < 0.0001$). However, despite comparable performance in the task (salient target fovea vs. extrafovea: 92%$\pm$2 vs. 94%$\pm$3, Wilcoxon rank-sum test: $p = 0.4$, Cohen's $d = -0.73$), we found that the latency of microsaccades was longer compared to the latency of saccades when the stimuli were presented foveally for a salient target (fovea vs. extrafovea: 311ms$\pm$30ms vs. 232ms$\pm$30ms, Wilcoxon rank-sum test: $p = 0.005$, Cohen's $d = 2.4$). Interestingly, we did not observe any latency differences between saccades and microsaccades when the target was not salient (fovea vs. extrafovea: 422ms$\pm$80ms vs. 445ms$\pm$93ms, Wilcoxon rank-sum test: $p = 0.6$, Cohen's $d = -0.24$) in the absence of a salient distractor. Fovea vs. extrafovea: 429

ms ±76 ms vs. 440 ms ±90 ms, Wilcoxon rank-sum test: $p = 0.7$, Cohen's $d = -0.13$, in the presence of a salient distractor.

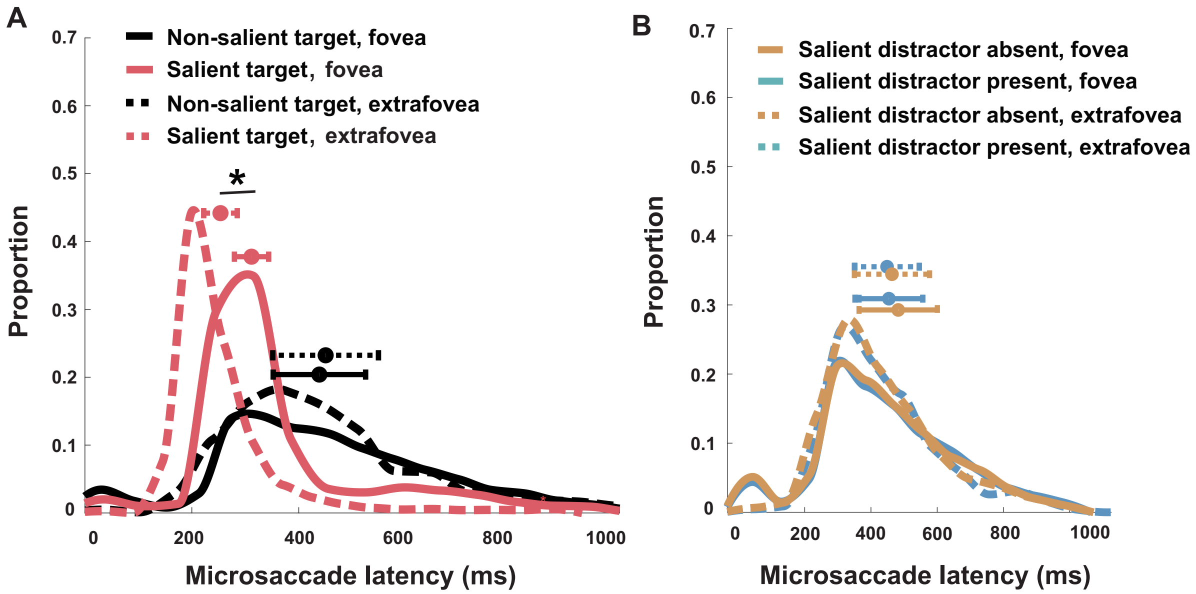


**Figure S1: Comparison between saccade and microsaccade latency in a foveal and extrafoveal search task.** $A$ - Average microsaccade latency for a salient target and non-salient target. Asterisks mark a statistically significant difference (Wilcoxon rank-sum test: $p < 0.0001$). $B$ - Average microsaccade latency in the presence and absence of a salient distractor. Dashed line indicate parafoveal condition. Solid line indicates foveal condition. Individual error bars on top mark the median latency.

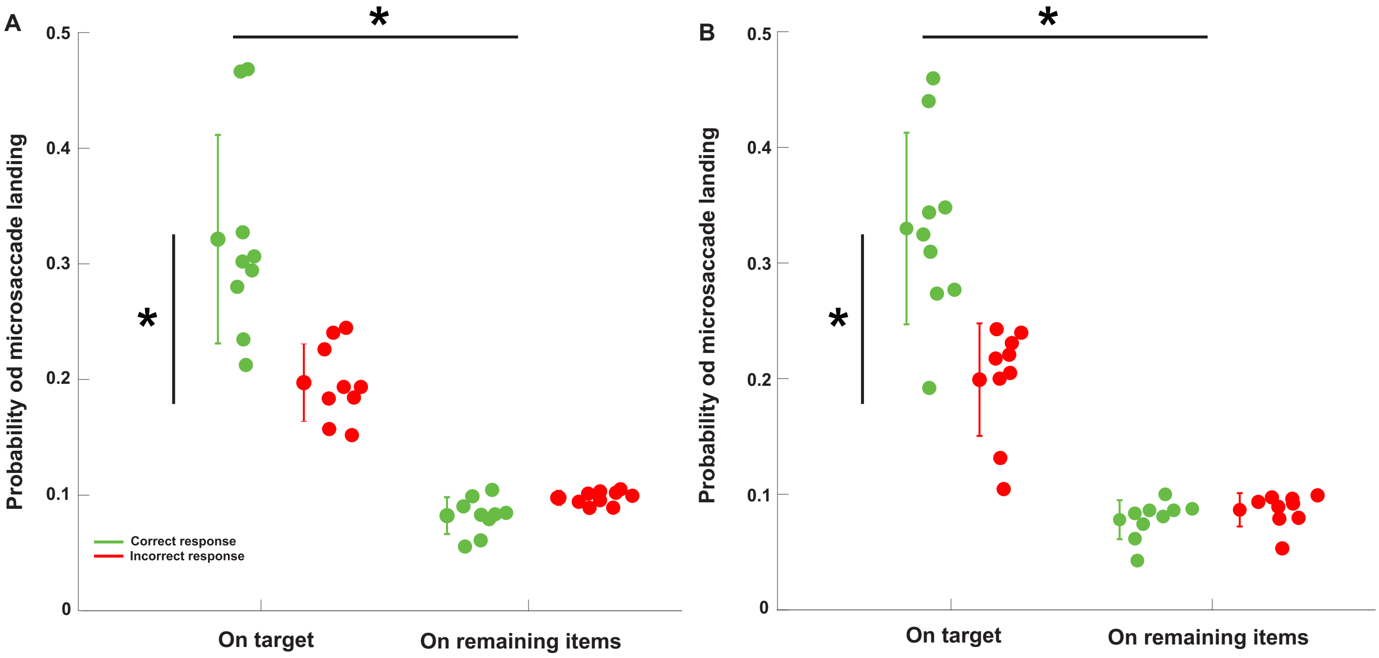


**Figure S2: Microsaccades accuracy in correct and incorrect trials.** Probability of microsaccade landing on the target and on remaining items in the stimulus array when a salient distractor was absent ($A$) and when it was present ($B$) in the stimulus array. Asterisks mark a statistically significant difference (two-tailed paied t-test: $p < 0.01$). Errorbars are SEM.
